# Nuclear DNA Damage Response Triggers Reorganization of Mitochondrial Nucleic Acids

**DOI:** 10.64898/2026.05.23.727397

**Authors:** Chenxiao Yu, Emma Evens, Sidney Tchiong, Marite Castromonte Albinagorta, Joshua Okletey, Marco Tigano

## Abstract

Mitochondria are subcellular organelles responsible for energy production, and a hub for several cellular signaling pathways that ultimately control cellular processes ranging from cell death to innate immunity. Genotoxic stress, including cellular irradiation, has been shown to cause the mitochondrial-dependent activation of innate immunity *via* release of mitochondrial nucleic acids in the cytosol. Yet, how the other cellular events triggered by genotoxic stress affects mitochondria and mitochondrial immunity is largely unexplored. Nuclear DNA damage responses (DDR) are a set of well-described responses to genotoxic stressors that allow the cells to, through characterized mechanisms including transcriptional responses and checkpoint activation, survive or die. Whether canonical DDR directly influence mitochondrial structure and activation of downstream pathways of immunity remains unclear. Here, we identify mito-blobs: enlarged TOMM20-positive mitochondrial structures induced by genotoxic stress that are enriched for TFAM-marked mitochondrial DNA nucleoids, FASTKD2-positive mitochondrial RNA granules, and immunogenic double-stranded RNA. While structurally resembling other mitochondria stress responsive rearrangements, mito-blobs carry the distinctive feature of being induced by nuclear DNA double-stranded breaks alone. Strikingly, mitochondrial double-stranded breaks failed to induce mito-blobs, indicating nuclear-to-mitochondrial signaling rather than an autonomous response to mitochondrial genome damage. Mechanistically, we show that mito-blobs formation strictly requires MFN1/2- and OPA1-dependent fusion machinery, while nuclear DNA damage invokes a classical ATM-p53 response, which lead to cell cycle block in the G1 phase that reduce DRP1 S616 phosphorylation and shifting mitochondrial morphology toward a low-fission pro-fusion state. Strikingly, the use of the standard of care CDK4/6 inhibitor palbociclib, was sufficient to trigger mito-blobs without nuclear DNA damage. Considering the mito-blobs’ high content in nucleic acids, we additionally investigated if affecting their life cycle could perturb inflammatory and interferon-associated gene expression downstream of genotoxic stress. We describe how autophagic-lysosomal clearance triggered upon cell cycle block limited mito-blob persistence, and blocking autophagy disposal unmasked a strong inflammatory response. Taken together, these findings suggest mito-blobs are the product of an active mitochondrial remodeling process elicited in response to nuclear genotoxic stress and that concentrate immunogenic mitochondrial nucleic acids in defined structures for their correct disposal via autophagy and avoid aberrant activation of innate immunity.

**Highlights:**

- Genotoxic stress induces enlarged mitochondrial structures called mito-blobs.
- Mito-blobs concentrate mitochondrial DNA, RNA granules, and double-stranded RNA.
- Nuclear DNA damage, but not mitochondrial, is sufficient to induce mito-blobs.
- CDK4 Inhibitors Trigger Mito-blob Formation.
- Mito-blob formation and clearance alter inflammatory gene expression.

**eTOC blurb:** Yu et al. identify mito-blobs as enlarged mitochondrial structures that form after genotoxic stress. They show that nuclear DNA breaks – as opposed to mitochondrial DNA breaks - trigger mito-blob formation through nuclear-to-mitochondrial signaling involving altered DRP1 phosphorylation and mitochondrial fusion machinery. Mito-blob persistence is limited by autophagic clearance and is associated with inflammatory and interferon-associated gene expression after stress.

## Introduction

Nuclear DNA damage is a central consequence of ionizing radiation and many genotoxic cancer therapies ^1^. DNA double-strand breaks activate the DNA damage response (DDR), including ATM-, ATR-, DNA-PK-, and p53-dependent signaling programs that coordinate DNA repair, checkpoint activation, cell-cycle arrest, senescence, cell death, and inflammatory gene expression ^2,3^. ATM-p53 signaling can induce p21/CDKN1A and suppress E2F- and CDK-associated cell-cycle programs, providing a well-established route from DNA damage to G1 arrest and reduced cell-cycle kinase activity^4^.

Mitochondria contain their own double-stranded circular genome of 16,6 Kbp which encodes for 13 of the >1,100 proteins in the mitochondrial proteome. The mitochondrial genome is packaged into dense proteinaceous structures rich in the transcription factor TFAM and called nucleoids^5,6^. Nucleoids are distributed throughout a dynamic mitochondrial network whose shape is controlled by fission and fusion pathways. DRP1-dependent fission and MFN1/2- and OPA1-dependent fusion regulate mitochondrial morphology ^7^, distribution, and quality control ^8^. Mitochondrial dynamics is also regulated throughout the cell cycle. The CDK1-cyclin B complex phosphorylates DRP1 at a conserved site DRP1 S616 (in humans), promoting mitochondrial fission ^9,10^ to prepare the cell for mitosis and ensure that mitochondria are correctly redistributed in the daughter cells. Given the intimate connection between mitochondrial dynamics and cell cycle progression, and between cell cycle arrest and DDR, it is tempting to speculate that DDR could influence mitochondrial dynamics and related responses. In support of this, both ATM and p53 can modulate mitochondrial functions, but independent of each other and independent of their canonical transcriptional programs ^11–14^.

Mitochondrial nucleic acids also have important signaling potential. Mitochondrial DNA, and double-stranded (mt-dsRNA) can promote inflammatory or interferon-associated responses when exposed to innate immune sensors ^15–17^ and have been implicated in the establishment of cellular senescence by activating the senescence associated secretory phenotype ^18–20^. However, how genotoxic stress influences the dynamic rearrangement of mitochondria to promote the release of nucleic acids and activation of downstream pathways is not well defined.

Prior studies have described enlarged mitochondrial nucleic acids-containing mitochondrial structures, including “mitobulbs”^21^, or mitochondrial RNA stress bodies^22^, particularly in settings of impaired mitochondrial fission or altered RNA processing ^23–26^. Here, we identify mito-blobs as enlarged TOMM20-positive mitochondrial structures induced by genotoxic stress and enriched for TFAM-marked nucleoids, FASTKD2-positive mitochondrial RNA granules, and J2-positive dsRNA signal. We show that nuclear DNA damage, but not mitochondrial, is sufficient to induce mito-blob formation. Mechanistically, our data support a model in which ATM-p53 signaling reduces DRP1 S616 phosphorylation and suppresses mitochondrial fission, thereby promoting mitochondrial remodeling into mito-blobs, a process supported by mitochondrial fusion. In parallel, we show that autophagic-lysosomal clearance of mito-blob contents is required to protect cells from innate immunity activation. Together, these findings define mito-blobs as a regulated mitochondrial remodeling state that links nuclear DNA damage signaling to mitochondrial dynamics and to inflammatory/interferon-associated gene expression.

## Results

### Genotoxic stress induces rearrangement of mitochondrial matrix contents and mitochondrial nucleic acids

We sought to investigate whether cancer cells treated with ionizing radiation (IR) experience mitochondrial nucleic acid stress. Lung adenocarcinoma A549 cells were cultured on glass coverslips and fixed to assess mitochondrial network morphology by immunofluorescence with the mitochondrial marker TOMM20. Mitochondrial nucleoids were counterstained with a primary antibody against TFAM. Under basal conditions (0 Gy), A549 cells exhibited a perinuclear, fragmented mitochondrial network (Fig. 1A), as reported in the literature ^27^.

**Figure 1.**
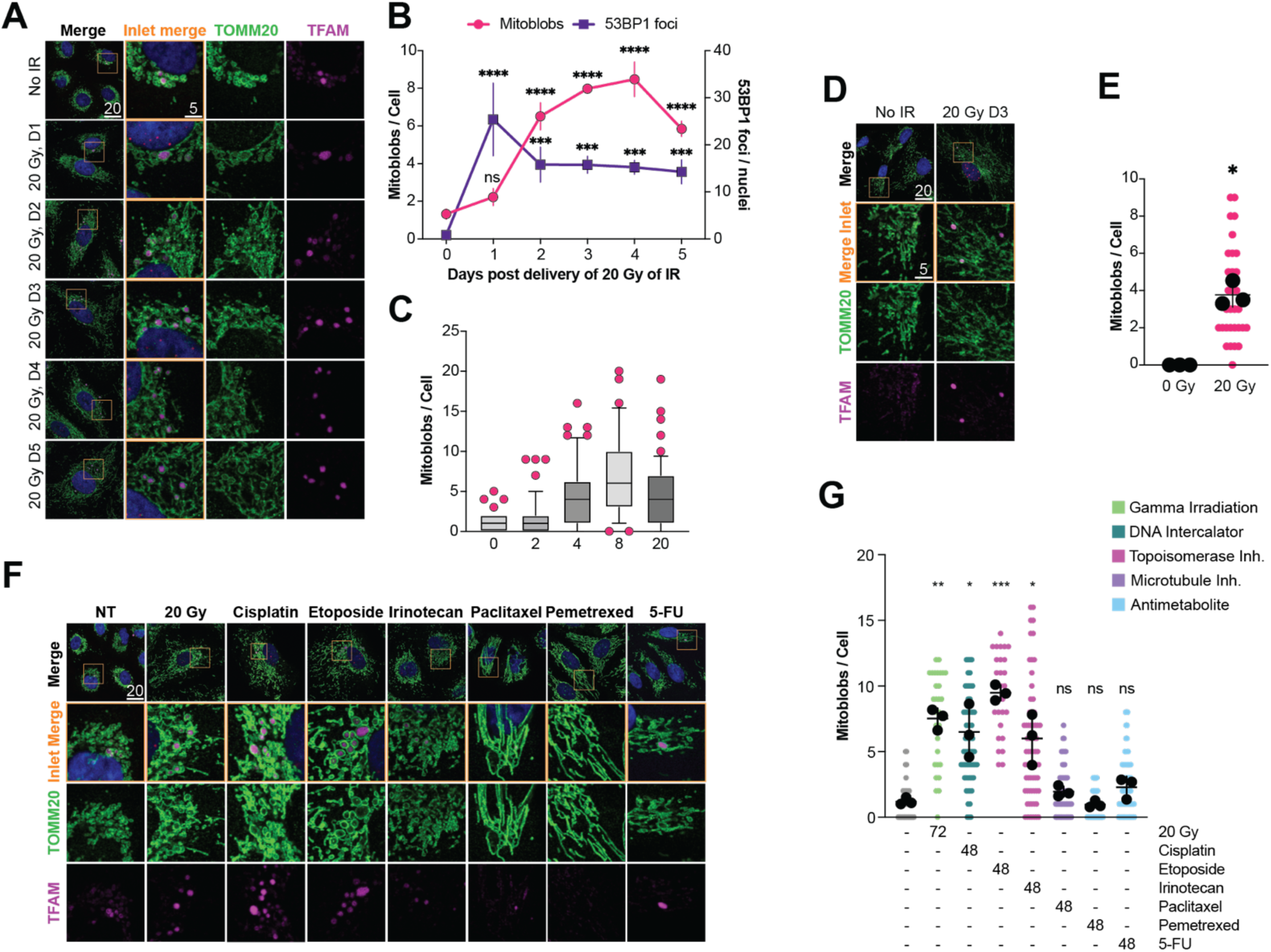
Genotoxic stress induces mito-blob accumulation. (A) Representative immunofluorescence images of A549 cells stained for TOMM20 and TFAM at the indicated times after 20 Gy ionizing radiation (IR). Enlarged TOMM20-positive mitochondrial structures containing concentrated TFAM signal are defined as mito-blobs. Scale bars, 20 µm; insets, 5 µm. (B) Quantification of mito-blobs per cell and 53BP1 foci per nucleus over time after 20 Gy IR. (C) Quantification of mito-blobs per cell 3 days after the indicated doses of IR. (D) Representative immunofluorescence images of ARPE-19 cells stained for TOMM20 and TFAM under untreated conditions or 3 days after 20 Gy IR. Scale bars, 20 µm; insets, 5 µm. (E) Quantification of mito-blobs per cell in ARPE-19 cells under the conditions shown in (D). (F) Representative immunofluorescence images of A549 cells stained for TOMM20 and TFAM after treatment with IR or the indicated cytotoxic agents. Drug-treated cells were treated with 100 nM cisplatin, etoposide, irinotecan, paclitaxel, pemetrexed, or 5-FU for 48 h. Scale bars, 20 µm; insets, 5 µm. (G) Quantification of mito-blobs per cell after the treatments shown in (F). IR was quantified 72 h after 20 Gy; drug-treated cells were quantified after 48 h treatment. Data are shown as individual cells with biological replicate means and mean +/- SEM or as box-and-whisker plots where indicated. n = 3 biological replicates except in (C), where one representative experiment is shown. ns, not significant; *p<0.05; **P < 0.01; ***P < 0.001; ****P < 0.0001 by 2 tailed Welch’s t test.

Most mitochondrial nucleoids appeared as small puncta distributed throughout the mitochondrial network, although a subset of A549 cells also contained enlarged TFAM-positive mitochondrial structures (Fig. 1A). Upon delivery of 20 Gy of IR, A549 cells rapidly accumulated bright 53BP1 puncta in the nuclear compartment that were partially resolved 48 hours after irradiation (Fig. S1A), indicative of nuclear double-stranded breaks (nDSBs). The number of 53BP1 foci did not return to baseline, consistent with published data ^28^. At the same time, we observed a time- and dose-dependent gradual accumulation of enlarged TOMM20-positive structures harboring a strong, diffuse TFAM signal (Fig. 1A–C), resembling enlarged nucleoid structures ^15^ — albeit of much larger size — or the condensed mitochondrial RNA granules that accumulate upon mitochondrial fission impairment ^25^.

Mitochondria with this morphology have previously been referred to as "blob" structures^29^; we will refer to them as mito-blobs hereinafter. Mito-blobs began to accumulate shortly after delivery of 20 Gy of IR — coincident with the appearance of 53BP1 foci — indicative of a response downstream of genotoxic stress (Fig. 1B). Mito-blobs reached a peak between days 3 and 4 post-irradiation before declining and persisted even after 53BP1 foci were partially resolved (Fig. 1B). When cells were irradiated at increasing doses (Fig. S1B), the number of mito-blobs increased in a dose-dependent manner up to 8 Gy, beyond which no further accumulation was observed (Fig. 1C and S1B). To determine whether mito-blob formation is cancer-specific or reflects a more general cellular response, we examined mito-blob formation in normal diploid ARPE-19 cells, a retinal pigment epithelial model. Ionizing radiation induced mito-blobs in ARPE-19 cells similarly to A549 (Fig. 1D–E and S1C–D), suggesting that mito-blobs represent a general response to genotoxic stress.

Given that ionizing radiation damages cells through multiple mechanisms — including single-and double-stranded DNA breaks and oxidative damage caused by water radiolysis ^30^ — we next asked whether mito-blob formation is specific to IR or reflects a broader response to genotoxic agents. A549 cells were treated with chemotherapy drugs spanning several classes: the DNA intercalator cisplatin, topoisomerase inhibitors etoposide and irinotecan, the microtubule inhibitor paclitaxel, and antimetabolites pemetrexed and 5-FU. Nuclear DNA damage was confirmed for each agent by 53BP1 immunofluorescence after 48 hours.

Cisplatin, etoposide, and irinotecan all produced robust 53BP1 accumulation and mito-blob induction at levels comparable to, or exceeding, those of IR (Fig. 1F–G). By contrast, paclitaxel, pemetrexed, and 5-FU caused minimal 53BP1 accumulation (Fig. S1E–F) and no significant mito-blob induction (Fig. 1F–G). These data indicated that mito-blob formation tracked with the ability of an agent to produce nuclear DNA damage, rather than with cytotoxicity per se. Next, we characterized the composition of mito-blobs.

### Mito-blobs are enlarged mitochondrial structures enriched for mtDNA, mitochondrial RNA granules, and J2-positive dsRNA

The mitochondrial gene expression machinery is organized in closely related processing hubs within the mitochondrial matrix ^31^, including nucleoids and RNA granules^32^. Several cellular stressors have been reported to dynamically reshape these machineries and their associated nucleic acids^22,25^. We therefore asked whether mito-blobs are enriched for mtDNA and mt-dsRNA — an immunogenic byproduct of mitochondrial transcription — and whether their composition changes upon IR. To mark mito-blobs we used TOMM20 to identify mitochondrial structures and antibodies against TFAM or FASTKD2 (an RNA granule marker) to define mito-blob boundaries. MtDNA was stained with an anti-dsDNA antibody, and mt-dsRNA was detected with the J2 antibody. Using confocal imaging, we detected all signals in A549 cells (Fig. 2A, D, G), including mt-dsRNA, which accumulates at variable levels in lung cancer cell lines ^33^. Line scan analysis confirmed that both mtDNA and mt-dsRNA are present in mito-blobs under basal conditions (Fig. 2B, E, H). Relative to the TOMM20 signal, TFAM and anti-dsDNA signal tended to cluster centrally, with dsDNA in a punctate distribution; FASTKD2 and mt-dsRNA accumulated more peripherally, closer to the TOMM20 boundary. After IR, mt-dsRNA showed the most pronounced peripheral shift (Fig. 2G–H). The resolution limits of current confocal imaging precluded confident quantification of this distribution, and super-resolution approaches will be needed to characterize the internal architecture of mito-blobs.

**Figure 2.**
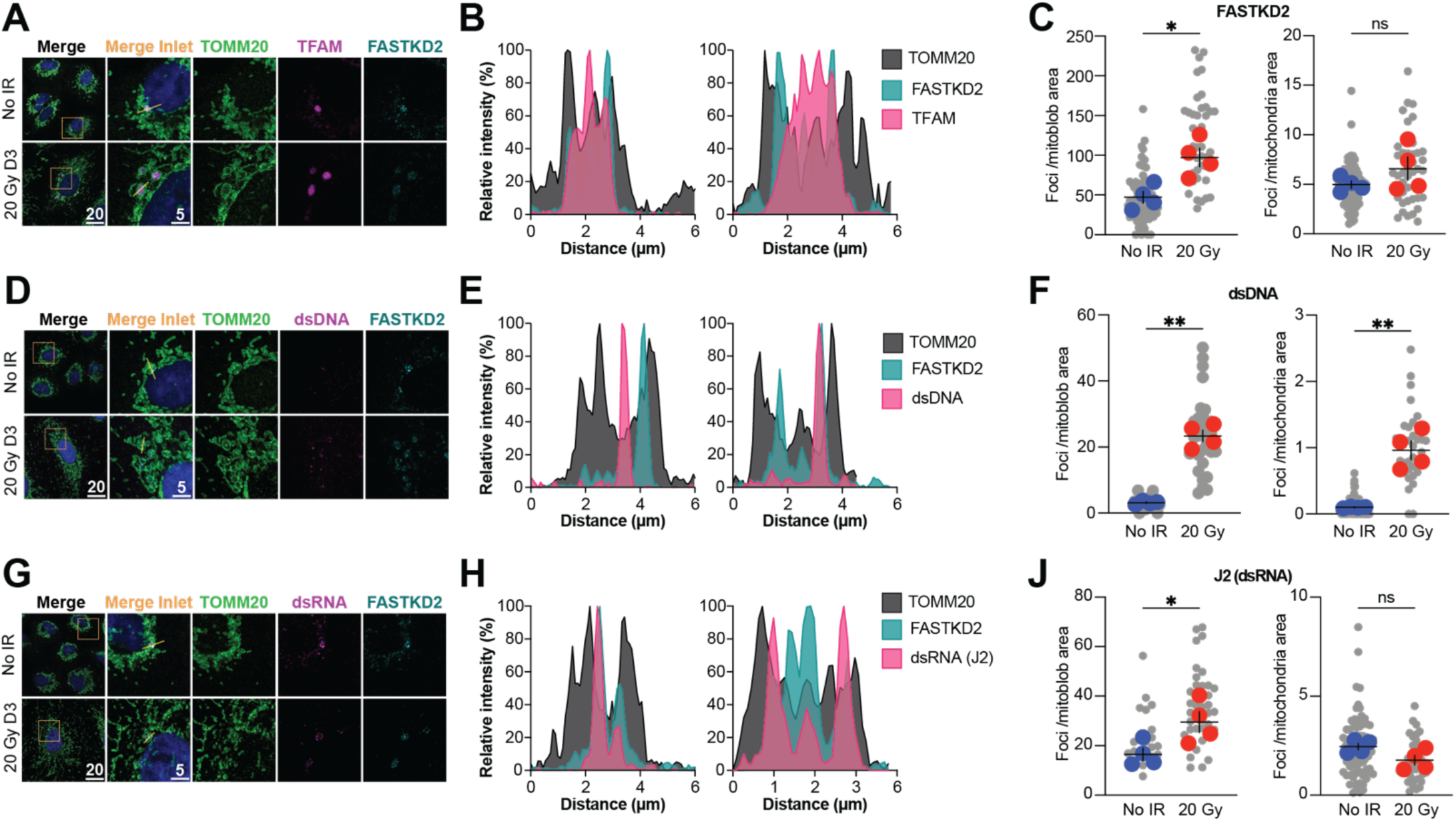
Mito-blobs are enriched for mitochondrial RNA granule marker FASTKD2, anti-DNA signal, and J2-positive dsRNA signal. (A) Representative immunofluorescence images of A549 cells stained for TOMM20, TFAM, and FASTKD2 under untreated conditions or 3 days after 20 Gy IR. Scale bars, 20 µm; insets, 5 µm. (B) Line-scan analysis of TOMM20, TFAM, and FASTKD2 relative fluorescence intensity across the regions indicated in (A). (C) Quantification of FASTKD2 foci density within mito-blob regions and across the total mitochondrial area under untreated conditions or 3 days after 20 Gy IR. (D) Representative immunofluorescence images of A549 cells stained for TOMM20, anti-DNA/dsDNA signal, and FASTKD2 under untreated conditions or 3 days after 20 Gy IR. Scale bars, 20 µm; insets, 5 µm. (E) Line-scan analysis of TOMM20, FASTKD2, and anti-DNA/dsDNA relative fluorescence intensity across the regions indicated in (D). (F) Quantification of anti-DNA/dsDNA-positive foci density within mito-blob regions and across the total mitochondrial area under untreated conditions or 3 days after 20 Gy IR. (G) Representative immunofluorescence images of A549 cells stained for TOMM20, J2-positive dsRNA signal, and FASTKD2 under untreated conditions or 3 days after 20 Gy IR. Scale bars, 20 µm; insets, 5 µm. (H) Line-scan analysis of TOMM20, FASTKD2, and J2-positive dsRNA relative fluorescence intensity across the regions indicated in (G). (J) Quantification of J2-positive dsRNA foci density within mito-blob regions and across the total mitochondrial area under untreated conditions or 3 days after 20 Gy IR. Data are shown as individual cells with biological replicate means and mean +/- SEM. n = 4 biological replicates. ns, not significant; *P < 0.05; **P < 0.01 by 2 tailed Welch’s t test.

To quantify how IR affects the overall content of mitochondrial nucleic acids, we used automated masking of mito-blob and total mitochondrial areas to count foci and measure density (Fig. S2A). Total mtDNA foci per cell increased upon IR (Fig. 2F and S2C), consistent with the reported increase in mtDNA copy number following IR ^28,34^. This increase persisted when normalized to total mitochondrial mass or mito-blob area (Fig. 2F), suggesting compensatory mtDNA synthesis rather than simple concentration within blobs. In contrast, the total number of FASTKD2 and J2-positive mt-dsRNA foci per cell did not increase upon IR (Fig. S2B and D), but their density within mito-blob areas increased significantly — for FASTKD2 (Fig. 2C) and mt-dsRNA (Fig. 2J) — with a reciprocal decrease in density throughout the rest of the mitochondrial network. These data were consistent with active shuttling of RNA granule components into mito-blobs rather than increased synthesis, resembling the ATAD3A-mediated nucleoid trafficking observed upon DRP1 inactivation ^24^.

Together, these data defined mito-blobs as enlarged mitochondrial structures enriched for nucleoid and RNA granule markers and immunogenic mt-dsRNA. While total mtDNA content increased after IR in line with previously reported mitochondrial mass expansion, mt-dsRNA and FASTKD2 appeared to be actively redistributed into mito-blobs. We next sought to define the mechanism by which IR drives mito-blob formation.

### IR induces mito-blob formation through mitochondrial dynamics

The mitochondrial network continuously remodels through fission and fusion (Fig 3A)^35^. Following IR, A549 mitochondria shifted toward a more elongated morphology — evidenced by increased aspect ratio of TOMM20-stained mitochondrial tracks (Fig. 3B) — suggesting net increased fusion/fission ratio. We hypothesized that this shift in mitochondrial dynamics was responsible for mito-blob formation. To test this, we used short-hairpin RNA (shRNA)-mediated silencing of the major fission and fusion effectors: MFN1, MFN2, OPA1, DRP1, MIEF1, and MIEF2. Knockdown efficacy was assessed functionally by measuring changes in mitochondrial aspect ratio or WB. Silencing fusion genes decreased the aspect ratio, indicating fragmentation (Fig. S3A); silencing DRP1, MIEF1, or MIEF2 increased the aspect ratio, indicating net fusion (Fig. S3A, C-D).

**Figure 3.**
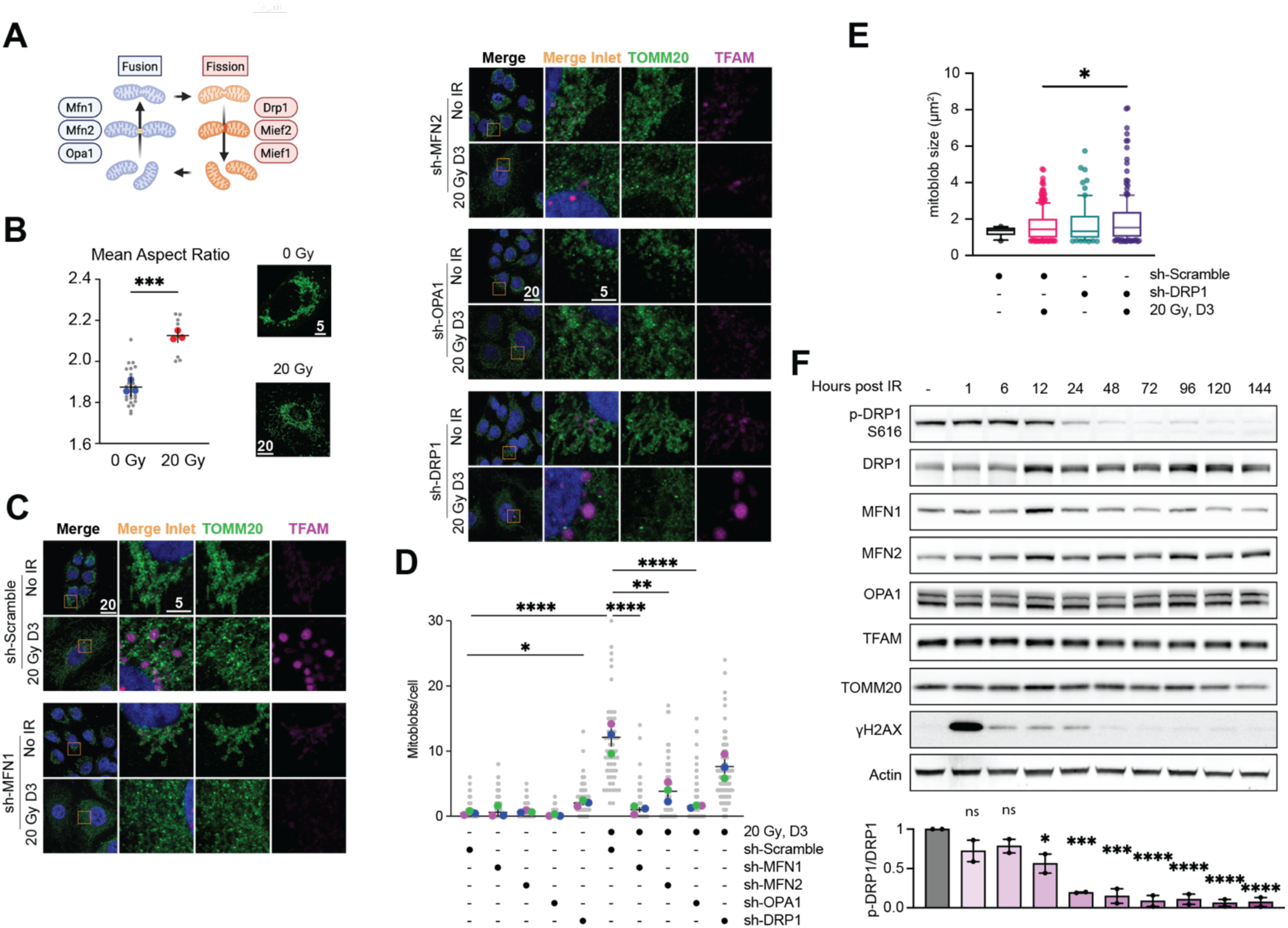
Mitochondrial dynamics and DRP1 S616 phosphorylation regulate IR-induced mito-blob formation and enlargement. (A) Schematic model of mitochondrial fission and fusion with main protein players. Created in BioRender. Tigano, M. (2026) https://BioRender.com/kwaybvd. (B) Representative TOMM20 images in A549 cells under untreated conditions or 3 days after 20 Gy IR, and quantification of mitochondrial aspect ratio (C) Representative immunofluorescence images of A549 cells expressing the indicated shRNAs under untreated conditions or 3 days after 20 Gy IR. Cells were stained for TOMM20 and TFAM to visualize mitochondrial morphology and TFAM-enriched mito-blobs. Scale bars, 20 µm; insets, 5 µm. (D) Quantification of mitoblobs per cell in A549 cells expressing sh-Scramble, sh-MFN1, sh-MFN2, sh-OPA1, or sh-DRP1 under untreated conditions or 3 days after 20 Gy IR. (E) Quantification of mito-blob size in sh-Scramble and sh-DRP1 cells under untreated conditions or 3 days after 20 Gy IR. (F) Immunoblot analysis of mitochondrial dynamics proteins, mito-blob-associated markers, and DNA damage response markers over time after 20 Gy IR. Intensity of p-DRP1 S616 is normalized to total DRP1 and then to no-IR group. Data are shown as individual cells with biological replicate means and mean +/- SEM, or as box-and-whisker plots where indicated. n = 3 biological replicates except in (E), where one representative experiment is shown. ns, not significant; *P < 0.05; ***P < 0.001 by 2 tailed Welch’s t test.

DRP1 knockdown alone caused a modest but significant increase in basal mito-blob number (Fig. 3C–D), indicating that reduced DRP1-dependent fission created a permissive state for mito-blob formation. However, knockdown of the DRP1 adaptors MIEF1 or MIEF2 did not affect basal mito-blob levels (Fig. S3C-D), supporting a specific role for DRP1 itself.

Knockdown of MFN1, MFN2, or OPA1 did not alter basal mito-blob number. Upon delivery of 20 Gy of IR, scramble-transduced cells accumulated a significant number of mito-blobs (Fig. 3D). This was unaffected by MIEF1 or MIEF2 knockdown (Fig. S3D). DRP1 knockdown did not further increase mito-blob number in IR conditions; instead, it caused a slight, non-significant decrease in number while significantly increasing mito-blob size (Fig. 3D-E). By contrast, knockdown of MFN1, MFN2, or OPA1 severely impaired the accumulation of large, TFAM-enriched mito-blobs after IR (Fig. 3C–D). Collectively, these data indicated that DRP1-dependent fission primarily regulated mito-blob size in irradiated cells, while complete fusion of both inner and outer mitochondrial membranes was required for mito-blob accumulation.

To complement the shRNA data, we examined the abundance of core mitochondrial dynamics proteins after IR by western blotting (Fig. 3F). Phosphorylation of DRP1 at S616 — a modification associated with active DRP1-mediated fission ^10^— declined progressively after IR, while total DRP1, MFN1, MFN2, and OPA1 levels remained stable (Fig. 3F). The decline in p-DRP1(S616) inversely correlated with mito-blob accumulation over the same time course (Fig. 1B), consistent with reduced DRP1 fission activity as a contributor to mito-blob formation.

### Nuclear DNA double-stranded breaks are sufficient to induce mito-blob formation

The data above suggested that genotoxic stress rewired mitochondrial dynamics toward a pro-fusion state, driving mito-blob formation. However, IR and cisplatin damage both nuclear and mitochondrial DNA ^30,36^, making it impossible to determine the relevant compartment from pharmacological experiments alone. To resolve this, we selectively introduced DSBs into either the mitochondrial or nuclear genome and asked which was sufficient to trigger mito-blobs.

MtDNA DSBs are rapidly followed by mtDNA degradation ^37–39^. To deliver mtDNA DSBs, we expressed an inducible mitochondria-targeted ApaLI restriction endonuclease — which cleaves the MT-RNR1 locus encoding 12S rRNA ^40^ — alongside a catalytically dead control, using a doxycycline-inducible lentiviral system in A549 cells (Fig. 4A). Because ApaLI-active caused mtDNA depletion and loss of TFAM signal, which would interfere with TFAM-based mito-blob scoring, we instead monitored mitochondrial morphology by live-cell labelling with MitoTracker Deep Red, using SYBR Green I to confirm mtDNA depletion (Fig. 4B). Despite confirmed mtDNA loss, ApaLI-active did not elicit mito-blob formation after 72 hours of induction (Fig. 4C). To confirm that ApaLI-transduced cells retained the capacity to remodel their mitochondrial network, we delivered 20 Gy of IR and observed the expected mito-blob accumulation. These results indicate that mtDNA damage is not a primary trigger of mito-blob formation under these conditions.

**Figure 4.**
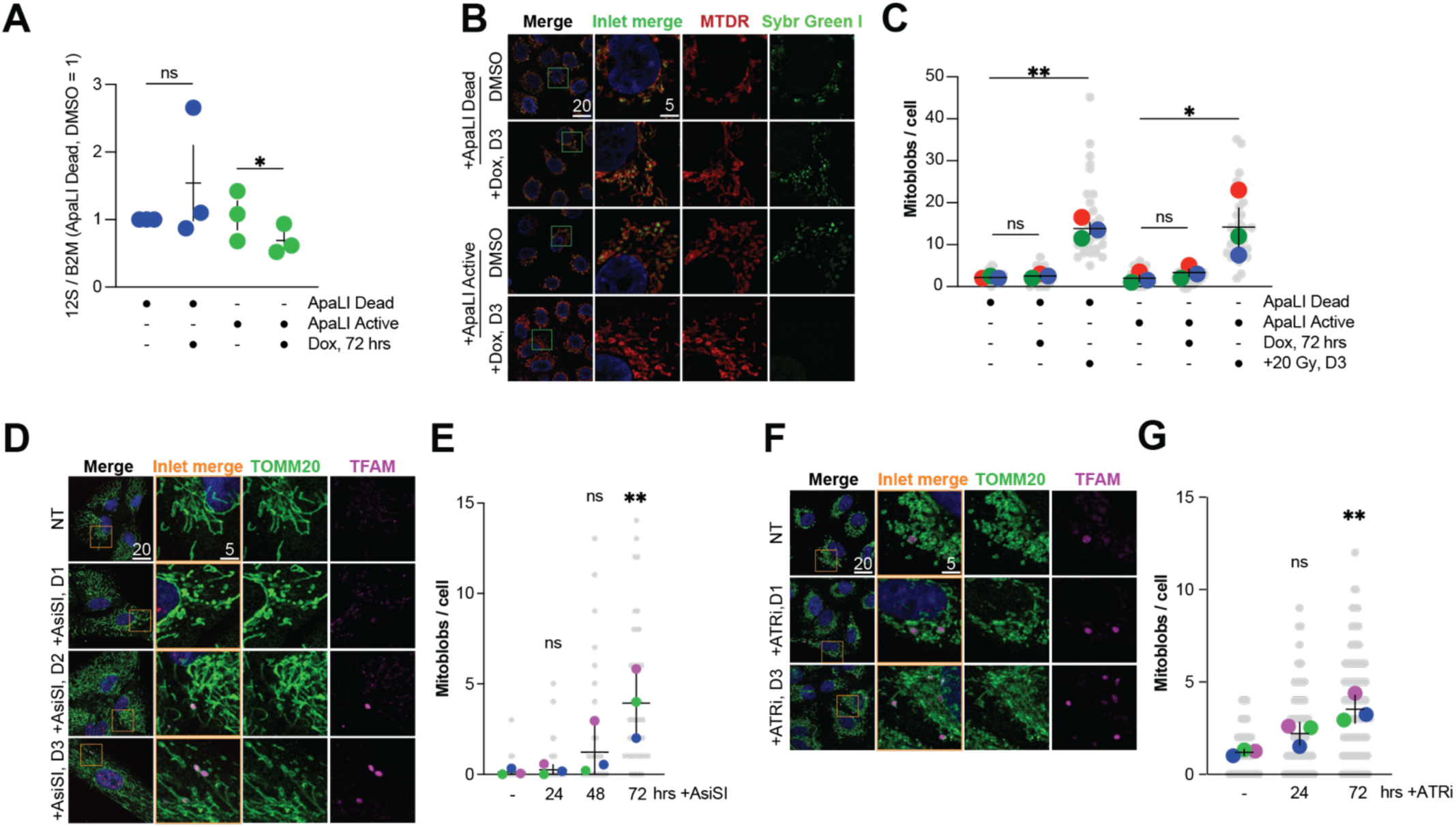
Mitochondrial and nuclear DNA perturbation assays for mito-blob formation. (A) Quantification of mitochondrial 12S rRNA locus abundance normalized to B2M in A549 cells expressing doxycycline-inducible catalytically inactive mito-ApaLI (ApaLI Dead) or active mito-ApaLI (ApaLI Active), with or without 72 h doxycycline treatment. (B) Representative live-cell images of A549 cells expressing ApaLI Dead or ApaLI Active after DMSO or 3 days doxycycline treatment. Cells were stained with MitoTracker Deep Red (MTDR) and SYBR Green I. Scale bars, 20 µm; insets, 5 µm. (C) Quantification of mito-blobs per cell in A549 cells expressing ApaLI Dead or ApaLI Active under the indicated doxycycline and IR conditions. Doxycycline was applied for 72 h where indicated; IR-treated cells were quantified 3 days after 20 Gy IR. (D) Representative immunofluorescence images of ARPE-19 cells carrying the inducible AsiSI system at the indicated times after AsiSI induction. Cells were stained for TOMM20 and TFAM. Scale bars, 20 µm; insets, 5 µm. (E) Quantification of mito-blobs per cell at the indicated times after AsiSI induction. (F) Representative immunofluorescence images of A549 cells treated with ATR inhibitor (ATRi) for the indicated times. Cells were stained for TOMM20 and TFAM. Scale bars, 20 µm; insets, 5 µm. (G) Quantification of mito-blobs per cell at the indicated times after ATRi treatment. Data are shown as individual cells or biological replicate values with biological replicate and means and mean +/- SEM. n = 3 biological replicates. ns, not significant; *P < 0.05; **P < 0.01 by 2 tailed Welch’s t test.

We next asked whether nuclear DSBs alone are sufficient. ARPE-19 cells were chosen for this experiment because they formed mito-blobs in response to IR (Fig. 1D–E) but did not accumulate them at baseline, providing a clean readout. These cells were transduced with an inducible AsiSI restriction enzyme system, which generates approximately 150 DSBs across the nuclear genome ^41^. AsiSI induction triggered robust nuclear DNA damage as confirmed by 53BP1 immunofluorescence (Fig. S4A–B). Strikingly, mito-blob number increased progressively after AsiSI induction, reaching significance by day 3 (Fig. 4D–E). To validate this result in A549 cells with an independent approach, we treated cells with an ATR inhibitor (ATRi), which induces nuclear DNA damage (Fig. S4D–E) without affecting mtDNA copy number (Fig. S4C; ddC was used as a positive control for mtDNA depletion). ATR inhibition also led to time-dependent mito-blob accumulation in A549 cells (Fig. 4F–G).

Together, these experiments established that nuclear DSBs were sufficient to induce mito-blob formation, while mtDNA damage or depletion was not. Our data therefore supported a model in which a nuclear-to-mitochondrial anterograde signaling pathway, triggered by nDSBs and acting in part via loss of p-DRP1(S616), drove the reorganization of the mitochondrial network into mito-blobs.

### Nuclear DSBs signal to mitochondria via the ATM–p53 axis to drive mito-blob formation

Our results implicate anterograde DDR-to-mitochondria signaling, with reduced p-DRP1(S616) as a candidate mechanistic link. To identify the relevant signaling pathway, we first performed unbiased RNA sequencing in A549 cells before and 3 days after 20 Gy IR. The transcriptome was substantially altered, with 883 genes upregulated and 1139 genes downregulated (Fig. 5A). Gene set enrichment analysis revealed downregulation of G2M checkpoint and E2F target gene sets, and upregulation of the p53 pathway and epithelial–mesenchymal transition signatures (Fig. 5A). Notably, CDK1 — a key regulator of DRP1 S616 phosphorylation — was among the most downregulated genes, consistent with G1 arrest downstream of ATM–p53–p21 signaling and with the loss of p-DRP1(S616) we observed after IR.

**Figure 5.**
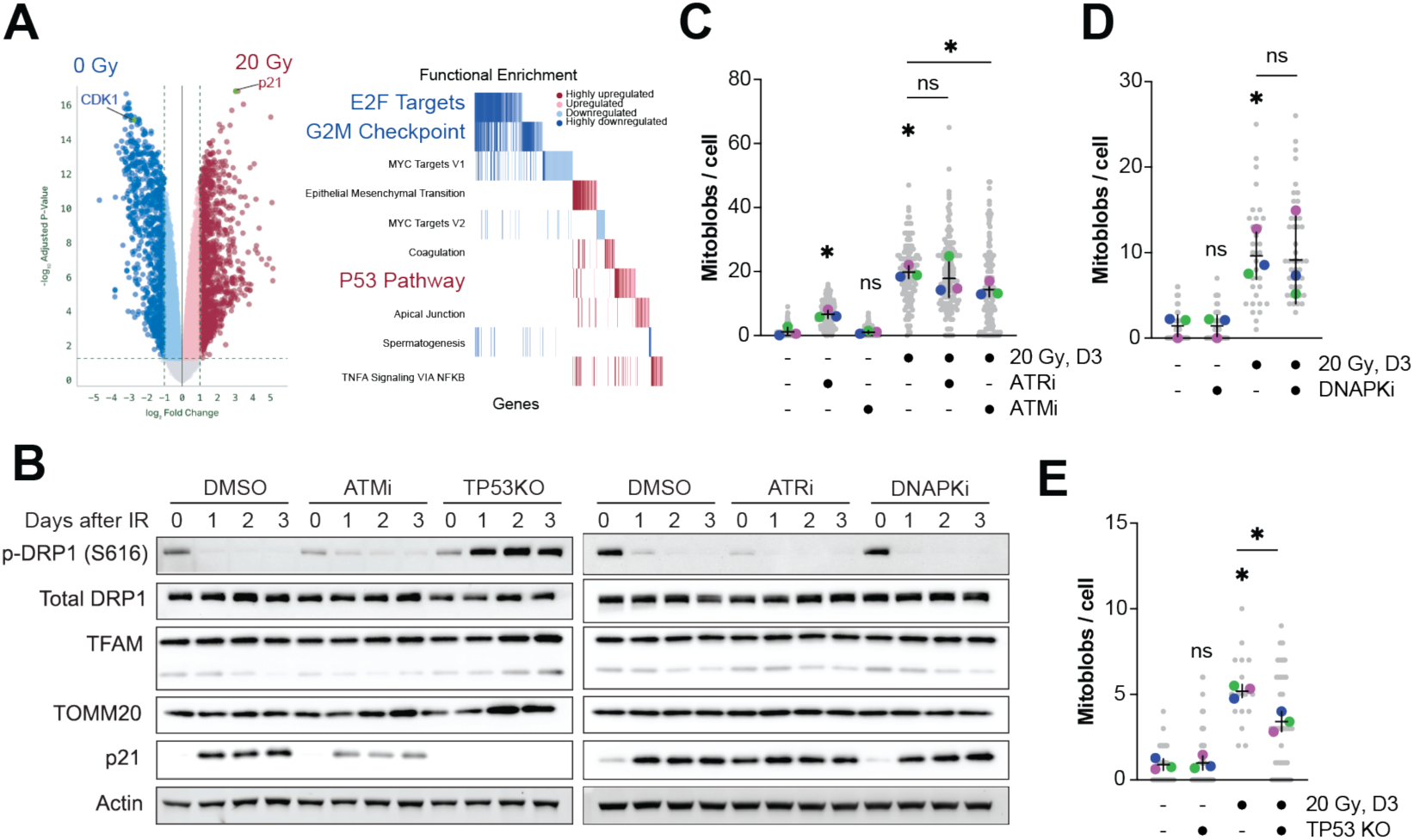
DDR kinase and TP53 perturbations during IR-induced mito-blob formation and DRP1 S616 phosphorylation. (A) RNA-seq analysis of A549 cells under untreated conditions or 3 days after 20 Gy IR. Left, volcano plot highlighting selected genes including CDK1 and p21/CDKN1A. Right, pathway-level functional enrichment summary of selected gene sets. (B) Immunoblot analysis of p-DRP1 S616, total DRP1, TFAM, TOMM20, p21, and actin in A549 cells under the indicated conditions over 0-3 days after 20 Gy IR. Conditions include DMSO, ATM inhibitor (ATMi), TP53 knockout (TP53KO), ATR inhibitor (ATRi), and DNA-PK inhibitor (DNAPKi). Actin was used as a loading control. (C) Quantification of mito-blobs per cell in A549 cells treated with DMSO, ATRi, or ATMi, with or without 20 Gy IR. IR-treated cells were quantified 3 days after irradiation. (D) Quantification of mito-blobs per cell in A549 cells treated with DMSO or DNAPKi, with or without 20 Gy IR. IR-treated cells were quantified 3 days after irradiation. (E) Quantification of large mito-blobs (diameter>1.44 µm (8 pixel)) per cell in WT and TP53KO A549 cells, with or without 20 Gy IR. IR-treated cells were quantified 3 days after irradiation. Data are shown as individual cells with biological replicate means and mean +/- SEM. n = 3 biological replicates except in (B), where representative immunoblots are shown. ns, not significant; *P < 0.05 by 2 tailed Welch’s t test.

To define which DDR kinases are required, we used CRISPR–Cas9 to generate TP53-knockout A549 cells (enriched by nutlin-3a selection; Fig. S5A) and pharmacological inhibitors of ATM (ATMi), ATR (ATRi), and DNA-PK (DNA-PKi). We monitored p-DRP1(S616) levels by western blotting for up to 3 days after IR (Fig. 5B, S5G-H). p-DRP1(S616) declined dramatically within 24 hours of IR; this decline was fully reversed by TP53KO and delayed by ATMi and ATRi, while DNA-PKi had no effect (Fig. 5B). Consistent with these effects on p-DRP1(S616), DNA-PKi did not alter IR-induced mito-blob accumulation, and ATRi had no significant effect on mito-blob number despite preserving p-DRP1(S616). ATMi, however, significantly reduced — though did not fully abolish — IR-induced mito-blobs (Fig. 5C-D).

TP53KO produced a more complex phenotype: in contrast to the pro-fusion shift seen in wild-type cells after IR (Fig. 3B), TP53KO cells maintained a fragmented mitochondrial network consistent with sustained high p-DRP1(S616) (Fig. S5A, E). This fragmented background complicated mito-blob scoring. When all mito-blob sizes were considered, no significant difference was detected between WT and TP53KO cells after IR (Fig. S5D). However, when analysis was restricted to larger mito-blobs (diameter>1.44 µm (8 pixel)), IR-induced accumulation was significantly reduced in TP53KO cells (Fig. 5E).

Immunoblot analysis further supports the ATM→p53 hierarchy. IR induced time-dependent p21 accumulation alongside p-DRP1(S616) loss (Fig. 5A–B). Albeit not significant, ATMi provoked a visible reduction in p21 induction and delayed p-DRP1(S616) loss, while TP53KO abolished p21 induction and restored p-DRP1(S616) after IR. Together, these data supported a model in which IR activated ATM, which signaled through p53 and p21 to repress CDK1 activity, thereby reducing p-DRP1(S616) and shifting the mitochondrial network toward the pro-fusion state required for mito-blob formation.

### Autophagic clearance of mito-blobs modulates innate immune gene expression

Because the ATM–p53–p21 axis drove G1 arrest, we asked whether G1 arrest alone — without nDSBs — was sufficient to induce mito-blobs. We treated A549 cells with the CDK4/6 inhibitor palbociclib ^42^ and monitored mito-blob number for up to 7 days. Palbociclib caused a detectable increase in mito-blob number from day 2 onward, peaking at day 3 (Fig. 6A). Unlike IR-induced mito-blobs, which persisted to at least day 5 (Fig. 1B), palbociclib-induced mito-blobs declined by day 5 and returned near baseline by day 7 (Fig. 6A). G1 arrest was therefore sufficient to induce mito-blobs, and the more rapid resolution compared with IR suggested that IR engaged additional mechanisms — potentially related to sustained DDR signaling or direct mitochondrial stress — that stabilized mito-blobs or slowed their clearance.

**Figure 6.**
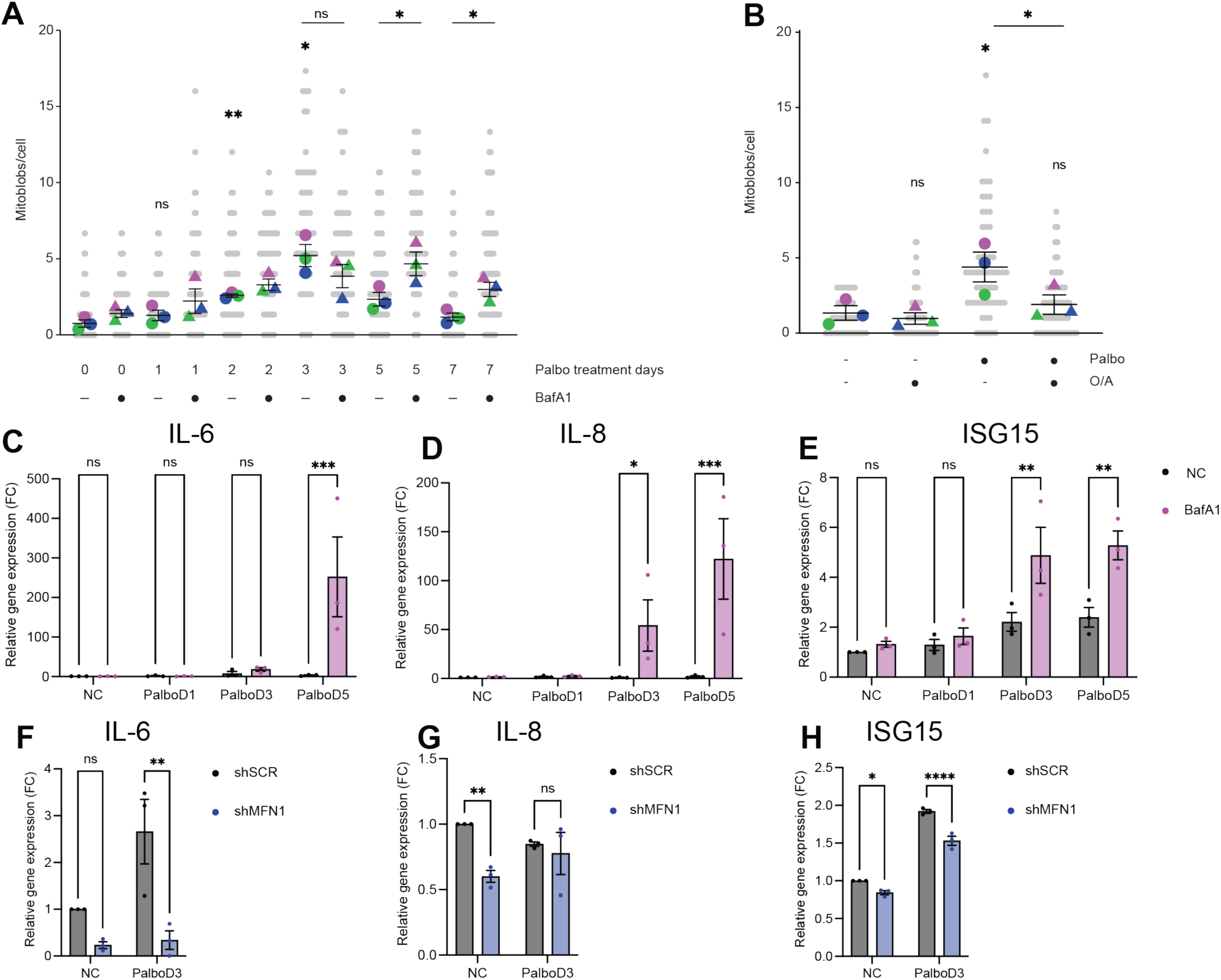
Mito-blob abundance and inflammatory/interferon-associated gene expression under palbociclib, lysosome inhibition, mitophagy induction, and MFN1 knockdown. (A) Quantification of mito-blobs per cell in A549 cells treated with palbociclib for the indicated number of days, with or without BafA1 treatment during the final 24 h before collection. (B) Quantification of mito-blobs per cell in A549 cells treated with palbociclib and/or oligomycin plus antimycin A (O/A). (C-E) RT-qPCR analysis of IL-6 (C), IL-8 (D), and ISG15 (E) expression in A549 cells under control conditions or after 1, 3, or 5 days of palbociclib treatment, with or without BafA1 during the final 24 h before collection. (F-H) RT-qPCR analysis of IL-6 (F), IL-8 (G), and ISG15 (H) expression in A549 cells expressing sh-Scramble or sh-MFN1 under control conditions or after 3 days of palbociclib treatment. Data are shown as individual cells or biological replicate values with mean +/- SEM. n = 3 biological replicates. ns, not significant; *P < 0.05; **P < 0.01; ***P < 0.001; ****P < 0.0001 by 2 tailed Welch’s t test (A, B) or two-way ANOVA (C-H).

The spontaneous resolution of palbociclib-induced mito-blobs prompted us to test whether autophagy mediated their clearance, consistent with the reports of p53-dependent mitophagy that occurred after IR independently of mitochondrial damage ^43^. We exposed palbociclib-treated cells to bafilomycin A1 (BafA1), a V-ATPase inhibitor that blocks lysosomal acidification, for the final 24 hours of each experimental time point (applied briefly at each time point due to toxicity). BafA1 did not significantly alter mito-blob numbers during the accumulation phase (days 1–3) but significantly increased them at days 5 and 7 (Fig. 6A and S6A), precisely when mito-blobs were normally cleared. This temporal specificity indicated that autophagy mediated mito-blob clearance rather than formation. Stimulating mitophagy with oligomycin plus antimycin A (O/A) — a Complex V/III inhibitor combination that activates mitophagy ^44^— had the opposite effect: when combined with palbociclib, O/A significantly reduced mito-blob accumulation compared with palbociclib alone, while O/A alone had no significant effect (Fig. 6B and S6B). Together, these data showed that mito-blobs were dynamically regulated by autophagic flux: lysosomal inhibition delayed clearance and mitophagy stimulation accelerated it.

To determine whether mito-blob clearance influences innate immune gene expression, we used BafA1 to prevent clearance in palbociclib-treated A549 cells and measured expression of IL-6, IL-8, and ISG15 by RT-qPCR at intervals over 5 days. Palbociclib alone did not induce ISG expression in A549 cells. However, the combination of palbociclib and BafA1 significantly induced all three ISGs (Fig. 6C-E), demonstrating that persistence of mito-blobs — rather than their formation per se — drove innate immune activation. To ask whether mito-blob formation is required for this response, we used MFN1 knockdown, which prevented mito-blob enlargement while leaving the baseline mitochondrial network intact (Fig 3C–D).

MFN1 knockdown reduced baseline ISG expression in unchallenged A549 cells (significantly for IL-6 and ISG15, not IL-8) and attenuated the ISG response under palbociclib (Fig. 6F-H). These two results — BafA1 increasing ISGs when clearance is blocked, and shMFN1 decreasing ISGs when formation is prevented — were complementary: both formation and clearance failure were required for full innate immune activation and disrupting either was sufficient to suppress ISG induction. These findings supported a model in which mito-blobs served as transient containers for immunogenic mitochondrial nucleic acids — particularly mt-dsRNA — that were degraded by autophagy to prevent innate immune activation, and implicated mito-blob-mediated sequestration as a mechanism by which cancer cells may limit inflammatory responses to genotoxic therapy.

## Discussion

The central finding of this study is the identification of mito-blobs – enlarged, TOMM20-positive mitochondrial structures induced by genotoxic stress that concentrate mitochondrial nucleic acids before being cleared by autophagy to limit innate immune activation. Morphologically anomalous mitochondrial structures have been described previously under different names and in different contexts. Mitochondrial "blobs" were first noted in stressed cells without functional characterization^29^. Mitobulbs – enlarged structures enriched for mtDNA – have been described in settings of impaired DRP1-dependent fission, including upon genetic inactivation of DRP1 itself ^24,25,45,46^. Most recently, mesoscale matrix bodies enriched in mt-dsRNA, termed mitochondrial RNA stress bodies (MSBs), were identified in response to RNA processing stress and linked to translational repression ^22^. Mito-blobs share features with all three: they are enlarged mitochondria, enriched for nucleoid markers, and contain mt-dsRNA. However, they are uniquely defined by a specific upstream trigger, nuclear DNA damage signaling, rather than intrinsic mitochondrial dysfunction. Critically, unlike mitobulbs, impaired fission alone does not drive mito-blob formation downstream of genotoxic stress. Similarly, unlike MSBs, the mt-dsRNA enrichment in mito-blobs reflects active shuttling of pre-existing RNA molecules rather than *de novo* synthesis or accumulation due to diminished degradation. Yet, we cannot exclude the possibility that mito-blobs, mitobulbs, and MSBs are all manifestations of a shared, stress-responsive mitochondrial reorganization program activated by distinct upstream signals.

The spatial organization of nucleic acids within mito-blobs suggests internal compartmentalization. TFAM fills the mito-blob volume broadly, whereas dsDNA signal concentrates centrally in discrete *puncta*. FASTKD2 and mt-dsRNA localize more peripherally, in proximity to the outer mitochondrial membrane. This arrangement becomes more pronounced after ionizing radiation, particularly for mt-dsRNA. The re-distribution could position immunogenic mt-dsRNA close to the outer membrane where cytosolic sensors are accessible but to substantiate this claim with sufficient confidence super-resolution approaches (STED or SIM) will be needed. Overall, total mtDNA increases throughout the mitochondrial network following IR, which is consistent with new mtDNA synthesis reported by others ^28,34^. Therefore, in contrast to mitobulbs generated in response to DRP1 inactivation, mtDNA is not actively recruited inside mito-blobs. By contrast, the total number of FASTKD2 and J2-positive mt-dsRNA *foci* is unchanged after IR, but their density increases specifically within the mito-blob area with a reciprocal decrease in the remainder of the network. This pattern is consistent with active shuttling of RNA granule components into mito-blobs, analogous to the afore mentioned trafficking of mtDNA within mitobulbs, a processed mediated by ATAD3A ^24^. Whether ATAD3A contributes to RNA granule redistribution in mito-blobs remains to be tested.

A central and unexpected finding is that mito-blob formation is triggered specifically by nuclear DNA damage and not by mitochondrial DNA damage. Ionizing radiation, cisplatin, etoposide can damage both genomes simultaneously and introduce mitochondrial stress, making difficult to understand where the signal to generate mito-blobs originates. We resolved this by delivering targeted DSBs to each compartment independently. Inducible mitochondrial restriction endonuclease cleavage via ApaLI, which depletes mtDNA downstream of mtDNA double-stranded breaks ^39,38,37^, failed to generate mito-blobs despite mtDNA depletion. Conversely, nuclear DSBs delivered by the inducible AsiSI system ^41^ were sufficient to produce mitob-lobs in ARPE-19 cells with kinetics mirroring those of ionizing radiation. ATR inhibition independently confirmed this in A549 cells. These experiments establish that the relevant signal originates in the nucleus. It is worth noting that mtDNA cleavage has its own well-characterized signaling outputs: in addition to mtDNA depletion-associated innate immune activation via mt-RNA sensing ^17^, mtDNA DSBs can trigger the integrated stress response through ATAD3A-dependent *cristae* destabilization ^40^. That these responses are engaged by mtDNA damage while mito-blob formation is not, illustrates that nuclear and mitochondrial DSBs activate distinct, non-redundant downstream programs.

This positions mito-blob formation as a new form of anterograde signaling from the DDR to the mitochondrial network. This is distinct from the two best-characterized pathways by which p53 and ATM reach mitochondria. Cytoplasmic p53 can translocate to the outer mitochondrial membrane to engage BCL-XL and BAX and trigger transcription-independent apoptosis, a process regulated by MDM2-mediated monoubiquitination and independent of ATM-dependent phosphorylation ^12–14^. ATM can also phosphorylate PINK1 to promote mitophagy independently of nuclear DNA damage ^11^. While the processes of mitochondrial fission and fusion have direct ties with both apoptosis and mitophagy, in the context of ATM and p53 mitochondrial dynamics processes were not implicated. Therefore, mito-blobs represent a different mitochondrial reshaping outcome of DDR signaling.

The signaling route from nuclear DSBs to mitochondrial dynamics centers on the canonical ATM–p53–CDK1 axis. CDK1 phosphorylates DRP1 at S616, a modification required for active DRP1-mediated fission during mitosis ^10^. Following ionizing radiation, p-DRP1(S616) declines rapidly in an ATM- and p53-dependent manner. ATMi delays but does not abolish this decline, while TP53 knockout fully restores p-DRP1(S616) levels and prevents mito-blob accumulation. Further, transcriptomic profiling after irradiation, and immunoblotting underscore activation of p21 and repression of CDK1, which is predicted to lead to p-DRP1(S616) loss, consistent with mito-blob formation. ATMi and TP53KO cause the same level of decrease in mito-blob number, supportive of the two of them working in the same pathway. Yet, at this stage, we cannot exclude other independent contributions of ATM and p53 to mitochondria – including the ATM–PINK1 mitophagy axis or the transcription-independent mitochondrial functions of p53. Combinatorial genetic experiments would be required to fully separate their roles. Nevertheless, the transcriptomic, immunoblotting, and mito-blob data are most consistent with the canonical ATM–p53transcriptional axis.

From a mechanistic perspective, the shRNA screen of mitochondrial dynamics factors highlights distinct contributions from fission and fusion machineries. Knockdown of MFN1, MFN2, or OPA1 each abolishes mito-blob accumulation after IR, indicating that complete fusion of both outer and inner mitochondrial membranes is required. DRP1 knockdown, by contrast, modulates mito-blob size without substantially altering their number. This data is compatible with mitochondrial fission acting as a sizing mechanism, and fusion being a core initiating event. The protein levels of MFN1, MFN2, and OPA1 remain stable after IR without detectable changes in OPA1 long/short isoform ratio, a known regulator of inner membrane fusion activity ^47^. This suggests that similarly to Drp1, post-translational regulation might be involved. Future studies will involve probing phosphorylation and ubiquitination status of MFN1 and MFN2 ^48–50^.

The model connecting G1 arrest to p-DRP1(S616) loss predicts that G1 arrest itself, independent of DNA damage, should be sufficient to induce mito-blobs. To induce G1 arrest independent of nuclear damage, we treated A549 cells with the CDK4/6 inhibitor palbociclib. Palbociclib produced mito-blobs with an accumulation kinetics similar to that of ionizing radiation, underscoring that the CDK1-DRP1 could be the functional convergence point. This finding carries clinical weight as CDK4/6 inhibitors are being widely tested in the cancer field, and are the current standard of care in HR+ breast cancer. Yet a mitochondrial remodeling response to these agents has not been previously described. Relevant in this context, CDK4 loss in triple-negative breast cancer cells was recently shown to produce impaired mitochondrial fission and "giant mitochondria" with elevated mtDNA content ^51^, a phenotype consistent with mito-blobs. It should be noted that CDK4/6 inhibitors can also have independent off-target effects on mitochondrial metabolism and future work will be needed to understand the different contributions ^52^.

A notable difference between palbociclib and ionizing radiation is the persistence of the induced mito-blobs. Palbociclib-induced mito-blobs are quickly resolved by Day 7, while IR-induced mito-blobs seems to persist. This can be explained by the fact that IR might be engaging additional pathways influencing mito-blobs. Although direct mtDNA damage did not cause mito-blob formation, our group recorded extensive transcriptional responses triggered by irradiation of mitochondrial DNA null cells ^17^. The identify of such pathways may warrant reinterpretation in light of this mito-blob life cycle.

The resolution of palbociclib-induced mito-blobs prompted us to examine the role of autophagy and lysosomal degradation in mito-blob clearance. Bafilomycin A1 significantly increased mito-blob numbers only during the clearance phase of palbociclib treatment (after day 5) suggesting that autophagy is responsible for clearance rather than formation. We also predicted that stimulating mitophagy with oligomycin and antimycin A would stimulate mito-blobs clearance. Intriguingly, these findings are consistent with a role for p53-dependent induction of mitophagy after IR, independent of mitochondrial damage ^43^. The mitophagy adaptor SPATA18, a p53 transcriptional target was the main candidate in that case and could be a likely mediator of mito-blob recognition by the autophagic machinery. In an effort to identify the possible adaptors bridging mito-blobs to autophagic machinery, future experiments should target SPATA18 and additional canonical and non-canonical pathways^53^.

The functional importance of mito-blob clearance is revealed by its consequences for innate immune gene expression, which we tested with two key experiments. First, blocking lysosomal clearance with BafA1 in palbociclib-treated A549 cells robustly induced IL-6, IL-8, and ISG15 while palbociclib alone had no effects. This result is consistent with the reported effects of both general autophagy and mitophagy in limiting innate immune activation driven by cytosolic mitochondrial DNA. Disrupting these processes has been shown to enhance anti-tumor immune responses *in vivo* ^54,55^. Second, MFN1 knockdown, which is functionally required to make mito-blobs curb both baseline and palbociclib induced immunity. The reduction of baseline ISG expression by shMFN1 has to be interpreted in the light of the mito-blobs that already exist in unchallenged A549 and the fact that lung adenocarcinoma cell lines (including A549) harbor a constitutive readily detectable amount of mt-dsRNA ^33^. MFN1-dependent fusion is required to facilitate MAVS clustering on the mitochondrial outer membrane for antiviral RNA sensing ^56^.

The concerted action of mitochondrial fusion and autophagy disposal positions mito-blobs as a transient buffering compartment: immunogenic mt-dsRNA is actively concentrated in enlarged mitochondria, sequestered from cytosolic sensors, and then eliminated by lysosomal degradation. When genotoxic stress increases, cells might experience a restricted autophagic capacity which exposes the excess nucleic acids concentrated in the mito-blobs ultimately driving robust ISG expression. The mito-blob clearance pathway may therefore represent a mechanism by which cancer cells maintain immune quiescence following genotoxic therapy, with implications for the design of combination immunotherapy strategies.

In conclusion, this work defines an anterograde signaling circuit in which nuclear DNA damage instructs the mitochondrial dynamics machinery to concentrate immunogenic nucleic acids in mito-blobs for autophagic disposal. Mito-blobs act as a temporary containment structure, sequestering mt-dsRNA and mtDNA from cytosolic immune sensors before their lysosomal degradation. Critically, mitochondria do not activate this response autonomously, as direct damage to the mitochondrial DNA does not activate mito-blobs. Almost paradoxically, we reported that mtDNA drives immunity during genotoxic stress, suggesting that a balancing act between the buffering capacity of mito-blobs and the stress responses downstream of mtDNA damage are the ultimate decisive factor influencing the severity of innate immunity responses. Understanding this balance may help explain why certain tumors respond poorly to genotoxic therapy from an immunological standpoint, and whether disrupting mito-blob clearance could shift the immune landscape in favor of anti-tumor responses.

## Limitations of the study

Mito-blobs have been investigated in lung adenocarcinoma cell lines and ARPE-19 to provide a background with no base-line blobs. While structures were detected in both cell lines, future studies will need to involve multiple lung cancer cell lines and other types of cancers. The genotype of each cell line should also be considered, especially in terms of LKB1, KRAS and p16 mutations. Further, at the present time there is no evidence that mito-blobs could be operating *in vivo*. While mito-blobs have enough differentiating properties from previous stress structures (mitobulbs and MSBs), there is still a lack of clear definition. Resolving this will require direct comparative analysis, including super-resolution imaging and proteomic characterization. The identity of adapters bridging mito-blobs to the autophagic machinery have not been studied and their identity will be clarified in future analysis together with the identity of cytosolic sensors of immunity triggered when the autophagy machinery becomes overwhelmed.

## Acknowledgements

We are deeply grateful to Professor Agnel Sfeir at Memorial Sloan Kettering Cancer Center for two exceptionally insightful and meaningful discussions, which played a critical role in shaping the direction of this study. Without these discussions, the progress of this work would likely have been substantially delayed. We also thank Dr. Timo Rey from the MRC Mitochondrial Biology Unit, University of Cambridge, for developing and publishing the mtFociCounter FIJI plugin ^57^. This highly useful tool greatly facilitated our image analysis and was essential for the completion of this study. We further thank Dr. Gregory Mazo for developing and publishing the QuickFigures FIJI plugin ^58^, which greatly assisted us in image preparation during figure and manuscript preparation.

## Declaration of Generative AI and AI-assisted technologies in the writing process

During the preparation of this work, the authors used ChatGPT/Codex, Claude/Claude code and Edison to improve the readability, grammar, and clarity of the manuscript. The authors take full responsibility for the content of the publication.

## STAR methods

### Cell culture procedures and treatments

#### Cell culture procedures

All cells were maintained at 37C in a humidified incubator with 5% CO2. Human retinal pigment epithelial ARPE-19 cells (ATCC, CRL-2302) were immortalized with hTERT. The ARPE-19 cell line expressing AsiSIInd was obtained by transduction with a lentiviral construct, which was a gift from R. Greenberg, as described previously (citation). Human lung adenocarcinoma cell lines A549, human colorectal adenocarcinoma cell line DLD1, human cervical adenocarcinoma cell line Hela, ARPE-19 and 293T cells (ATCC, CRL-3216) used for lentiviral production were cultured in high glucose Dulbecco’s modified Eagle medium with GlutaMAX™ Supplement(DMEM; Gibco, 10566024) supplemented with 10% Fetal bovine serum (Gibco), 100 U/mL penicillin--streptomycin (Gibco), 0.1 mM MEM non-essential amino acids (Gibco), 1mM Sodium pyruvate (Gibco) and 50 μg/mL uridine (Sigma-Aldrich). Human lung adenocarcinoma H1299 and H23 are cultured in RPMI-1640 with the same supplement as DMEM.

### Treatments

Cells were subjected to genotoxic stress via exposure to X-ray irradiation (IR) (20 Gy, SSD 60cm at 150 cGy/mins) or pharmacological treatment with DNA-damaging chemotherapeutics, including platinum agents (Cisplatin), topoisomerase inhibitors (Etoposide, Irinotecan), and antimetabolites (5-Fluorouracil, Pemetrexed). Inhibitors including KU60019 (ATMi), Ceralasertib (ATRi), AZD-7648 (DNA-PKcsi), ASN007/Ravoxertinib (ERKi), Rapamycin (mTORC1i) and Palbociclib (CDK4/6i) were used. ddC was utilized to deplete mitochondrial DNA (mtDNA) and Bafilomycin A1 (BafA1) was used to inhibit the V-ATPase thus inhibiting the lysosome function. A full list of chemicals used in the study and relative concentration is available in Supplementary Table 1.

### Genome-wide RNA-sequencing and bioinformatics analysis

Total RNA was purified from A549 cells with the NucleoMag RNA kit (Macherey-Nagel) following the manufacturer’s instructions. Total RNA extracted from A549 cells was submitted to Plasmidsaurus for 3’ end-counting RNA-seq on the Illumina platform. Libraries were prepared by reverse transcription with a poly(dT)VN primer, followed by tagmentation and amplification with unique dual indices; UMIs were used for deduplication. Reads (∼90 bp, stranded) were trimmed with FastP v0.24.0, aligned to the human reference genome using STAR v2.7, and deduplicated with UMICollapse v1.1.0. Gene expression was quantified with featureCounts (subread v2.1.1) using strand-specific counting against 3’ UTR and exon features. Differential expression was assessed with edgePython v0.2.5, and functional enrichment was performed using GSEApy v0.12 with the MSigDB Hallmark gene set.

### RT–qPCR

Total RNA was purified using the NucleoMag RNA kit (Macherey-Nagel) according to the manufacturer’s instructions, which included Genomic DNA digestion. 300 nanograms of RNA were reverse-transcribed using the Maxima H Minus First Strand cDNA Synthesis Kit with dsDNase (Thermo Scientific). Quantitative PCR (qPCR) was subsequently performed for 40 cycles on an Applied Biosystems QuantStudio 5 Real-Time PCR System. Reactions were conducted in triplicates utilizing PowerUp™ SYBR™ Green Master Mix (Thermo scientific) in a total volume of 10 μl with standard cycling conditions. Relative gene expression was normalized using ACTB1 as a housekeeping gene, and all calculations were performed using the Quantstudio software. A list of the primers employed is provided in Supplementary Table 1. Splicing variants and exon junctions were retrieved from the Ensembl database (https://uswest.ensembl.org/index.html).

### Genomic DNA isolation and mtDNA copy number

Purified total genomic DNA was used for mtDNA copy number measurement by qPCR and southern blot. Cell pellets harvested were resuspended in a 400 ml PBS buffer containing 0.2% (w/v) SDS, 5 mM EDTA, and 0.2 mg/ml Proteinase K and incubated at 50 C for 6 hours with constant shaking at 1,000 rpm. DNA was precipitated by adding 0.3 M sodium acetate, pH5.2, and 600 ml isopropanol and kept at -20 C for over 2 hours. Precipitated DNA was centrifuged at 20,000 x g at four C for 30 minutes, followed by a 70% ethanol wash. DNA was then resuspended in the TE buffer (10 mM Tris-HCl, pH8.0, 0.1 mM EDTA) and quantified by Nanodrop. To determine the relative mtDNA copy number by qPCR, sequences specific to mtDNA (MT-RNR1/2) and the nuclear B2M locus (Table S1) were independently amplified from 25 ng of total genomic DNA using the same qPCR protocol described above. The Quantstudio software used the primary relative quantification method with mtDNA as the target and B2M as a reference.

### Western blot analysis

Cells were washed with cold PBS and then lysed with SDS lysis buffer (1% SDS, 10 mM TRIS and 1 mM EDTA) with Halt™ Protease and Phosphatase Inhibitor Cocktail (Thermo Scientific). Lysates were sonicated (Sonic Dismembrator Model 500, Fisher Scientific) for 10 cycles of 3 s on/3 s off at 15% amplitude. Lysates were clarified by centrifugation for 30 min at 14,800 rpm, 4 °C and the supernatant was quantified using the enhanced BCA protocol (Thermo Fisher Scientific, Pierce). Lysates were mixed with 4X laemmli loading buffer (SDS: 8.0%, Glycerol: 40.0%, β-mercaptoethanol: 8.0%, Bromophenol Blue: 0.02%, Tris-HCl: 250 mM) and then boiled at 70° C for 10 mins. An equal amount of proteins were loaded on SDS-PAGE gels and then transferred to a nitrocellulose membrane. Membranes were blocked in EveryBlot Blocking Buffer. Incubation with primary antibodies was performed overnight at 4 °C. Membranes were washed and incubated with HRP-conjugated secondary antibodies at 1:5,000 dilution, developed with Clarity ECL (Biorad) and acquired with a ChemiDoc MP apparatus (Biorad) and ImageLab v.5.2. Antibodies against actin were used as loading control dependent on the protein of interest. A full list of antibodies used in the study and relative dilutions is available in Supplementary Table 1.

### Immunofluorescence

Cells were plated on coverslips in 6-well plates and were treated with different chemicals and radiation for a certain time. culture media was removed and cells were fixed with PRE-WARMED 4% paraformaldehyde for 15 minutes at RT. After PBS washes, cells were permeabilized with 0.1% Triton X-100 in the PBS buffer for 5 minutes at RT, followed by PBS washes. Cells on coverslips were blocked with a blocking buffer (1%BSA+0.2%Gelatin+0.1%Triton X100) for 30-60 minutes at RT and incubated with primary antibody in the blocking buffer for 60 mins. Cells were then washed with PBS and hybridized with Alexa Fluor conjugated secondary antibody for 1 hour at RT. After PBS washes and DAPI staining, coverslips were mounted on glass slides with ProLong Gold Antifade mountant (Invitrogen). Slides were imaged on a Leica MICA/Stellaris microscope or Zeiss LSM880. Mitobulb was identified and masked via ilastik 1.4.1, and probability was saved as TIFF and then analyzed with CellProfiler. Briefly, objects are identified with the ‘IdentifyPrimaryObjects’ module with the Otsu algorithm and then objects larger than 1 um were included as mito-blob while larger than 1.5 µm as big mito-blob.

### Lentiviral delivery

shRNAs were cloned into a pLKO.1-Puro backbone as AgeI–EcoRI dsDNA oligomers (Integrated DNA Technologies) (Supplementary Table 1) and were introduced by four lentiviral infections at 12-h intervals in the presence of 8 μg ml−1 polybrene (Sigma-Aldrich) using supernatant from transfected HEK293T cells. pLKO.1-Puro was a gift from R. Weinberg (Addgene plasmid 845333). A549 and ARPE-19 cells were selected with 1 μg ml−1 puromycin for 5 days and recovered 1 additional day before evaluating the percentage of silencing. As experimental control, cells were infected with a pLKO.1-Puro plasmid encoding a scrambled shRNA sequence, which was a gift from D. Sabatini (Addgene plasmid 1864). Hairpin sequences were retrieved from the Broad Institute siRNA Consortium database (https://www.broadinstitute.org/rnai-consortium/rnai-consortium-shrna-library).

### CRISPR–Cas9 targeting

Polyclonal TP53-/- cell populations were generated as previously described ^59^. In brief, the CRISPR ribonucleoprotein (RNP) complex of guide RNA (gRNA) and Cas9 protein (IDT), were delivered into ARPE-19 and A549 cells using the Nucleofector system (Lonza). Cells that survived a treatment of 1μM idasanutlin were used for sanger sequencing and immunoblotting for p53 and p21.

### Statistical analysis

For RT-qPCR experiments, each biological replicate, sample, or gene was analyzed in technical triplicate. For imaging experiments, individual cells are displayed as gray dots to show cell-to-cell variability. When biological replicates were available, statistical tests were performed on biological replicate-level summary values rather than treating individual cells as independent biological replicates. Experiments quantified from a single independent experiment are explicitly indicated in the corresponding figure legends and are presented as representative or descriptive analyses rather than evidence of biological reproducibility.

Unless stated otherwise, statistical analyses were performed using 2-tailed Welch’s t tests in GraphPad Prism 11. Statistical significance was defined as P < 0.05 with a 95% confidence interval. Error bars and replicate numbers are defined in each figure legend.

---

**Figure S1.**
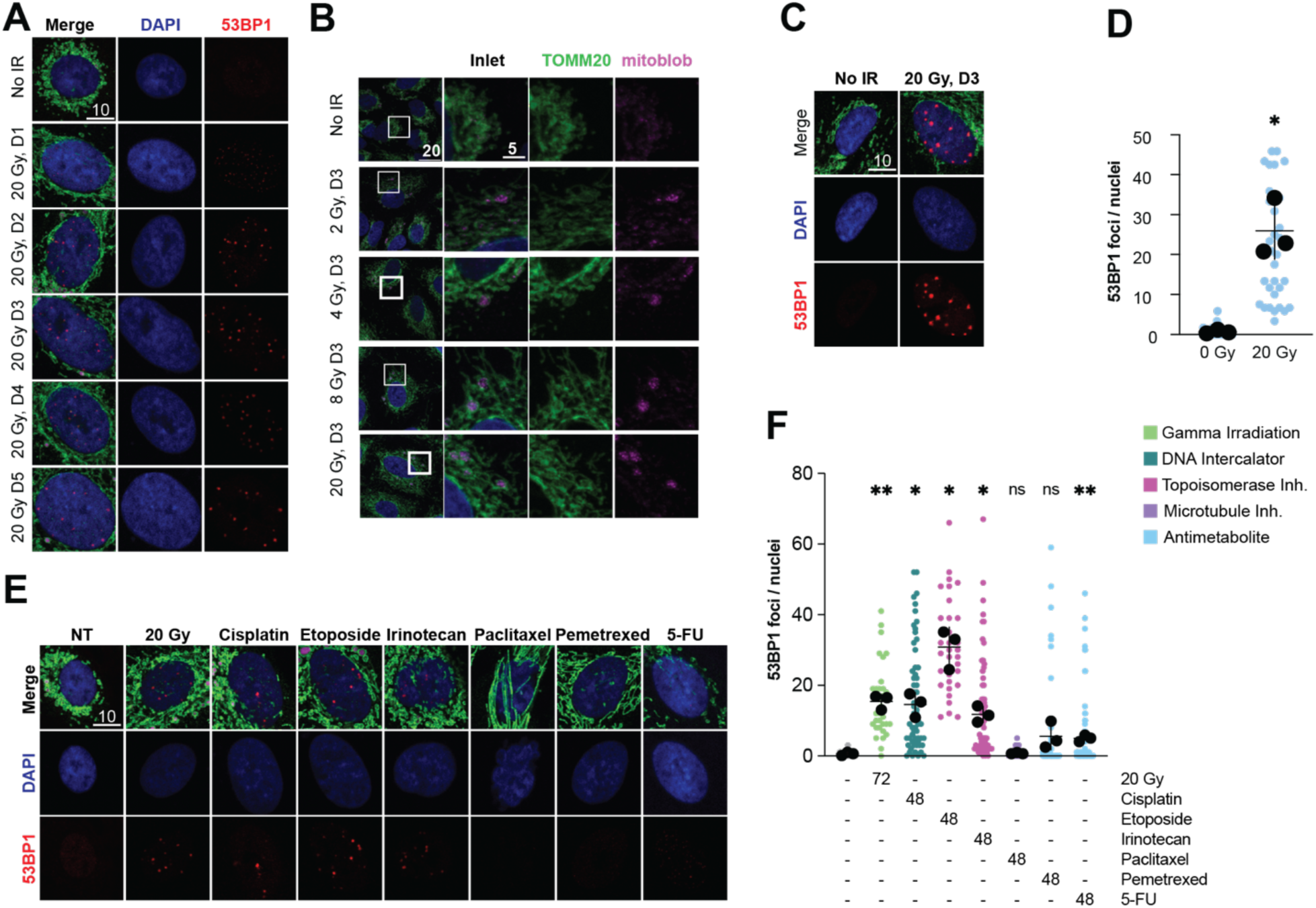
Nuclear DNA damage measurements and supporting mito-blob quantification. (A) Representative immunofluorescence images of A549 cells stained for DAPI and 53BP1 at the indicated times after 20 Gy IR. Scale bar, 10 µm. (B) Representative images showing TOMM20 staining and mito-blob (FASTKD2) in A549 cells 3 days after the indicated doses of IR. Scale bars, 20 µm; insets, 5 µm. (C) Representative immunofluorescence images of ARPE-19 cells stained for DAPI and 53BP1 under untreated conditions or 3 days after 20 Gy IR. Scale bar, 10 µm. (D) Quantification of 53BP1 foci per nucleus in ARPE-19 cells under the conditions shown in (C). (E) Representative immunofluorescence images of A549 cells stained for DAPI and 53BP1 after IR or the indicated cytotoxic treatments. Drug-treated cells were treated with 100 nM cisplatin, etoposide, irinotecan, paclitaxel, pemetrexed, or 5-FU for 48 h. Scale bar, 10 µm. (F) Quantification of 53BP1 foci per nucleus after the treatments shown in (E). IR was quantified 72 h after 20 Gy; drug-treated cells were quantified after 48 h treatment. Data are shown as individual nuclei with biological replicate means and mean +/- SEM. n = 3 biological replicates. ns, not significant; *P < 0.05; **P < 0.01; ***P < 0.001 by 2 tailed Welch’s t test.

**Figure S2.**
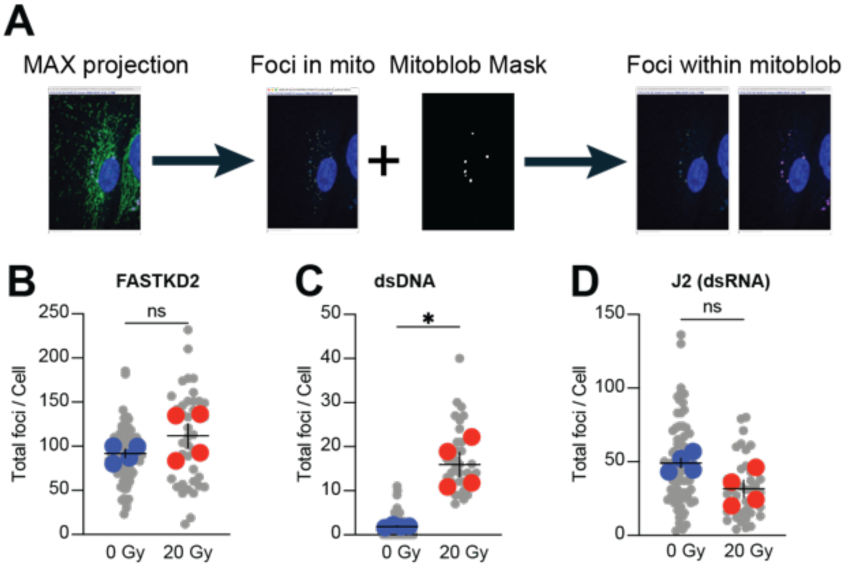
Workflow and total foci quantification supporting Figure 2 compartment analysis. (A) Schematic workflow for quantifying foci within mito-blob regions. Maximum-intensity projections were used to identify mitochondrial foci, mito-blob masks were generated independently, and foci within mito-blob regions were quantified from the overlap between mitochondrial foci and mito-blob masks. (B) Quantification of total mitochondrial FASTKD2 foci per cell under untreated conditions or 3 days after 20 Gy IR. (C) Quantification of total anti-DNA/dsDNA-positive mitochondrial foci per cell under untreated conditions or 3 days after 20 Gy IR. (D) Quantification of total J2-positive dsRNA mitochondrial foci per cell under untreated conditions or 3 days after 20 Gy IR. Data are shown as individual cells with biological replicate means and mean +/- SEM. n = 4 biological replicates. ns, not significant; *P < 0.05 by 2 tailed Welch’s t test.

**Figure S3.**
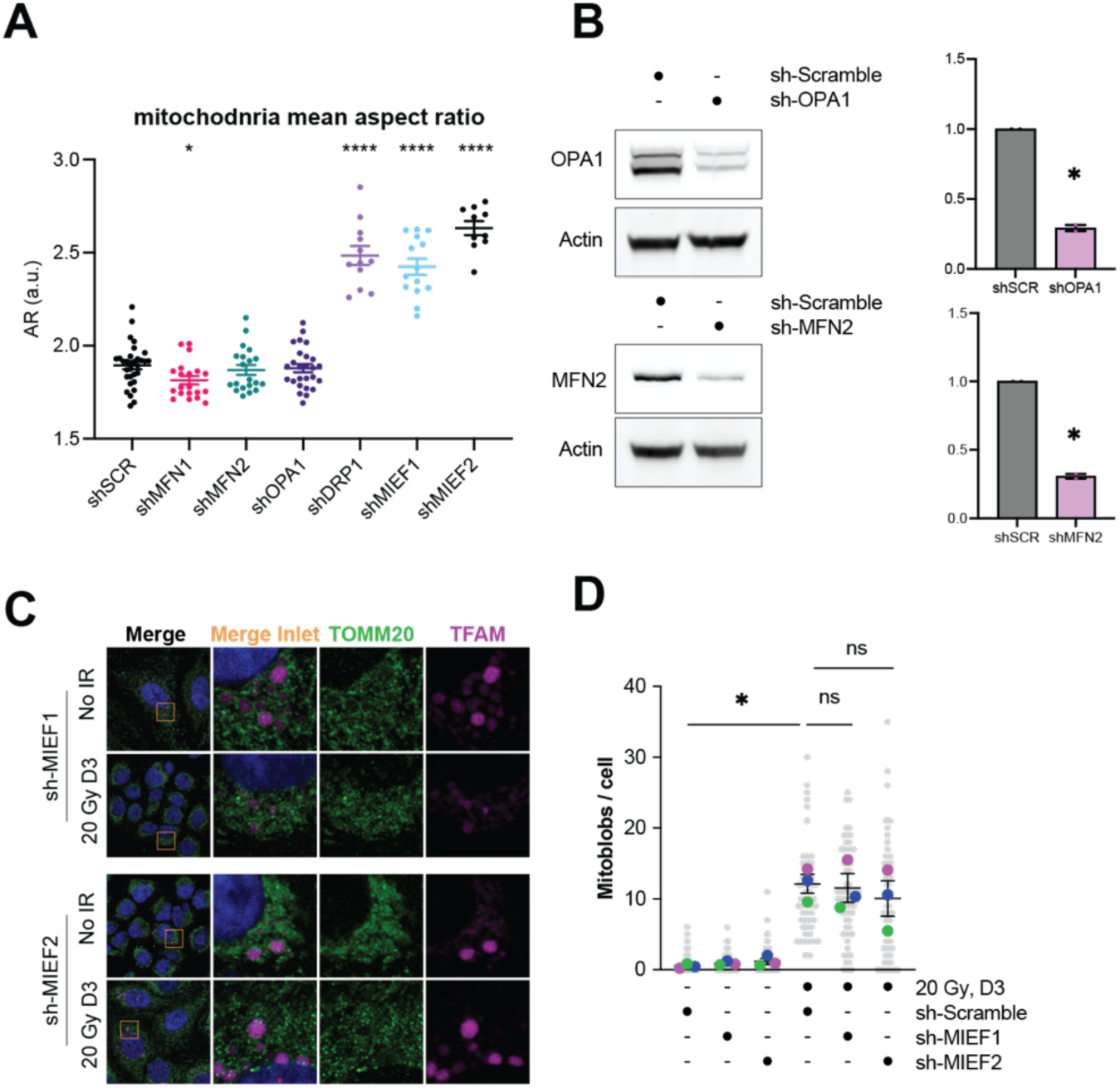
Supporting analysis of mitochondrial morphology and fission-factor knockdown. (A) Quantification of mitochondrial mean aspect ratio in A549 cells expressing sh-Scramble or shRNAs targeting MFN1, MFN2, OPA1, DRP1, MIEF1, or MIEF2. (B) Immunoblot analysis of OPA1 and MFN2 in A549 cells expressing the indicated shRNAs. Actin was used as a loading control. (C) Representative immunofluorescence images of A549 cells expressing sh-MIEF1 or sh-MIEF2 under untreated conditions or 3 days after 20 Gy IR. Cells were stained for TOMM20 and TFAM. Yellow boxes indicate regions shown in the insets. (D) Quantification of mito-blobs per cell in A549 cells expressing sh-Scramble, sh-MIEF1, or sh-MIEF2 under untreated conditions or 3 days after 20 Gy IR. Data are shown as individual cells with biological replicate means and mean +/- SEM. n = 4 biological replicates except in (A), where one representative experiment is shown. ns, not significant; *P < 0.05 ****P<0.0001 by 2 tailed Welch’s t test.

**Figure S4.**
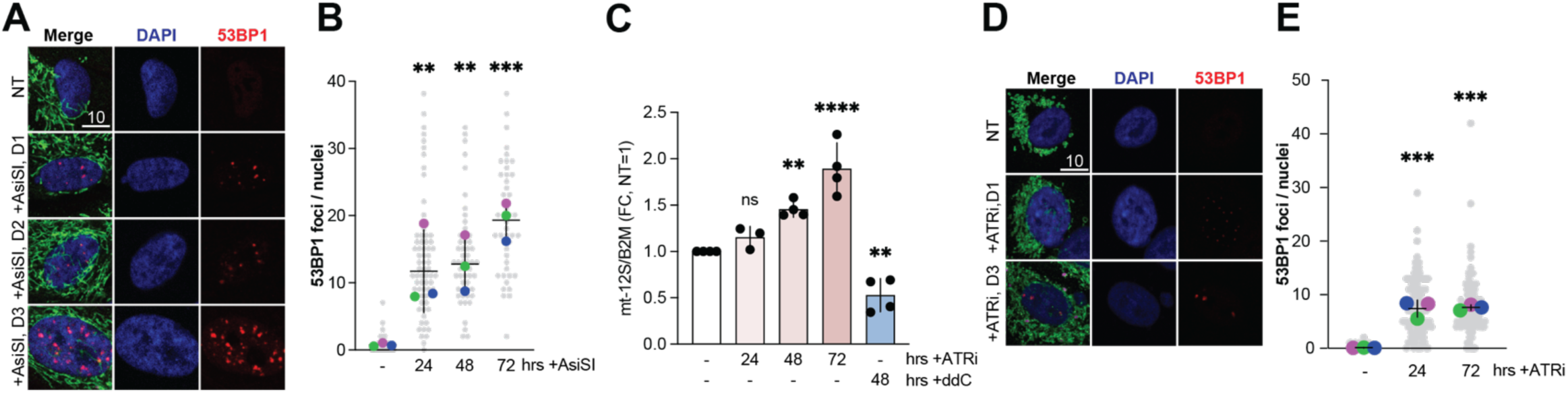
Nuclear DNA damage and mtDNA copy-number measurements supporting figure 4. (A) Representative immunofluorescence images of ARPE-19 cells carrying the inducible AsiSI system at the indicated times after AsiSI induction. Cells were stained for DAPI and 53BP1. Scale bar, 10 µm. (B) Quantification of 53BP1 foci per nucleus at the indicated times after AsiSI induction. (C) Quantification of mitochondrial 12S rRNA locus abundance normalized to B2M in A549 cells treated with ATRi for the indicated times or with ddC for 48 h as positive control for mtDNA depletion. (D) Representative immunofluorescence images of A549 cells treated with ATRi for the indicated times. Cells were stained for DAPI and 53BP1. Scale bar, 10 µm. (E) Quantification of 53BP1 foci per nucleus at the indicated times after ATRi treatment. Data are shown as individual nuclei or biological replicate values with biological replicate means and mean +/- SEM. n = 3 biological replicates. ns, not significant; **P < 0.01; ***P < 0.001; ****P < 0.0001 by 2 tailed Welch’s t test.

**Figure S5.**
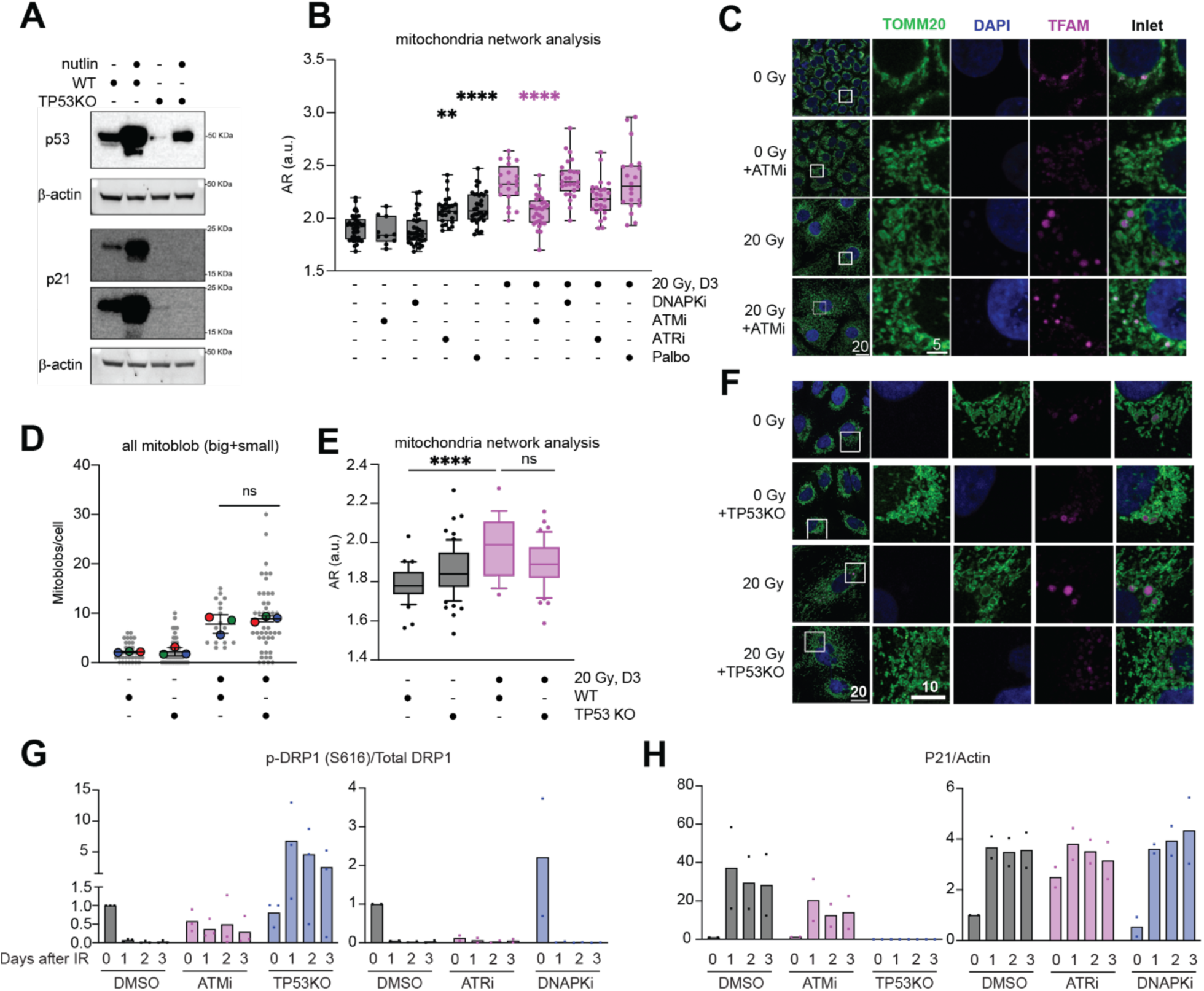
Supporting mitochondrial morphology and mito-blob measurements for Figure 5. (A) Immunoblot analysis of P53 and P21 in WT and TP53KO A549 before and after idasanutlin treatment. Actin was used as a loading control. (B) Quantification of mitochondrial network mean aspect ratio in A549 cells treated with the indicated inhibitors, with or without 20 Gy IR. IR-treated cells were quantified 3 days after irradiation. (C) Representative immunofluorescence images of A549 cells treated with DMSO or ATMi, with or without 20 Gy IR. Cells were stained for TOMM20, DAPI, and TFAM. Scale bars, 20 µm; insets, 5 µm. (D) Quantification of total mito-blobs per cell, including both large and small mito-blobs (diameter>0.9 um (5 pixels)), in WT and TP53KO A549 cells, with or without 20 Gy IR. n = 3 biological replicates. (E) Quantification of mitochondrial aspect ratio in WT and TP53KO A549 cells, with or without 20 Gy IR. (F) Representative immunofluorescence images of WT and TP53KO A549 cells, with or without 20 Gy IR. Cells were stained for TOMM20, DAPI, and TFAM. Scale bars, 20 µm; insets, 10 µm. (G-H) quantification of WB in Fig 5B. p-DRP1 is normalized to total DRP1 and p21 normalized to actin. Data are shown as means and mean +/- SEM or as box-and-whisker plots where indicated ns, not significant; **P < 0.01; ****P < 0.0001 by 2 tailed Welch’s t test.

**Figure S6.**
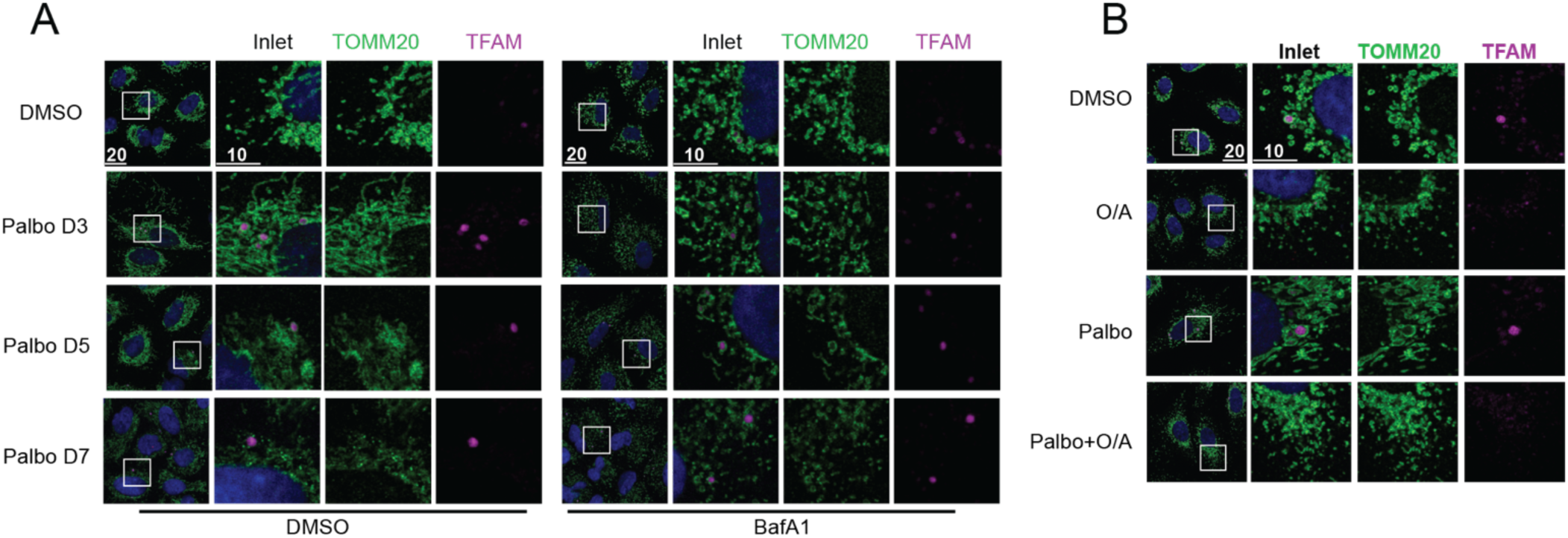
Representative images supporting Figure 6. (B) Representative immunofluorescence images of A549 cells treated with palbociclib for the indicated number of days, with or without BafA1 during the final 24 h before collection. Cells were stained for TOMM20 and TFAM. Scale bars, 20 µm; insets, 10 µm. (C) Representative immunofluorescence images of A549 cells treated with DMSO, O/A, palbociclib, or palbociclib plus O/A. Cells were stained for TOMM20 and TFAM. Scale bars, 20 µm; insets, 10 µm.

